# Metabolic alterations unravel the materno–fetal immune responses with disease severity in pregnant women infected with SARS-CoV-2

**DOI:** 10.1101/2023.05.10.540101

**Authors:** Sandhya Hora, Prabhjyoti Pahwa, Hamda Siddiqui, Anoushka Saxena, Minal Kashyap, Jayesh Kumar Sevak, Ravinder Singh, Maryam Javed, Pushpa Yadav, Pratibha Kale, Gayatri Ramakrishna, Asmita Rathore, Jaswinder Singh Maras, Shakun Tyagi, Shiv kumar Sarin, Nirupama Trehanpati

## Abstract

**Background:** Pregnancy being immune compromised state, COVID-19 disease poses high risk of premature delivery and threat to fetus. Plasma metabolome regulates immune cellular responses and we aimed to analyze the plasma secretome, metabolome and immune cells in COVID-19 positive pregnant mother and cord blood.

**Methods:** COVID-19 RT-PCR positive pregnant females (n=112) asymptomatic (n=82), or with mild (n=21) or moderate (n=9) disease and control healthy pregnant (n=10) females were included. Mother’s blood and cord blood (n=80) was analysed for untargeted metabolome profiling and plasma cytokines by high-resolution mass spectrometry (MS) and multiplex cytokine bead array. Immune scan in mothers was done using flow cytometry.

**Results:** In asymptomatic SARS-CoV-2 infection, --the amino acid metabolic pathways such as glycine, serine, L-lactate and threonine metabolism was upregulated, riboflavin and tyrosine metabolism, downregulated. In mild to moderate disease, the pyruvate and NAD^+^ metabolism (energy metabolic pathways) were mostly altered. In addition to raised TNF-α, IFN-α, IFN-γ, IL-6 cytokine storm, IL-9 was increased in both mothers and neonates. Pyruvate and NAD^+^ metabolic pathways along with IL-9 and IFN-γ had impact on non-classical monocytes, increased CD4 T cells and B cells but depleted CD8^+^ T cells. Cord blood mimicked the mother’s metabolomic profiles by showing altered valine, leucine, isoleucine, glycine, serine, threonine in asymptomatic and NAD^+^ and riboflavin metabolism in mild and moderate disease subjects.

**Conclusions:** Our results demonstrate a graduated immune-metabolomic interplay in mother and fetus in pregnant females with different degrees of severity of COVID-19 disease. IL-9 and IFN- γ regulated pyruvate, lactate TCA metabolism and riboflavin metabolism with context to disease severity are hall marks of this materno-fetal metabolome.

*Highlights:*

- SARS-CoV-2 infection alters energy consumption metabolic pathways during pregnancy.
- Pregnant women with mild to moderate COVID-19 show increased energy demands, and consume stored glucose by upregulating pyruvate and NAD^+^ metabolism.
- Increased TNF-α and IL-9 in mild COVID-19 disease involve TCA cycle to produce lactate and consume stored glucose by up regulating pyruvate and nicotinamide and nicotinate metabolism.
- With mild to moderate disease, raised IL-9 and TNF-α, decreased riboflavin pathway, exhaustion of T and B cells cause pathogenesis.
- Cord blood mimics the metabolic profile of mother’s peripheral blood, SARS-CoV-2 infection reshapes immune-metabolic profiles of mother-infant dyad.

*Graphical Abstract:* 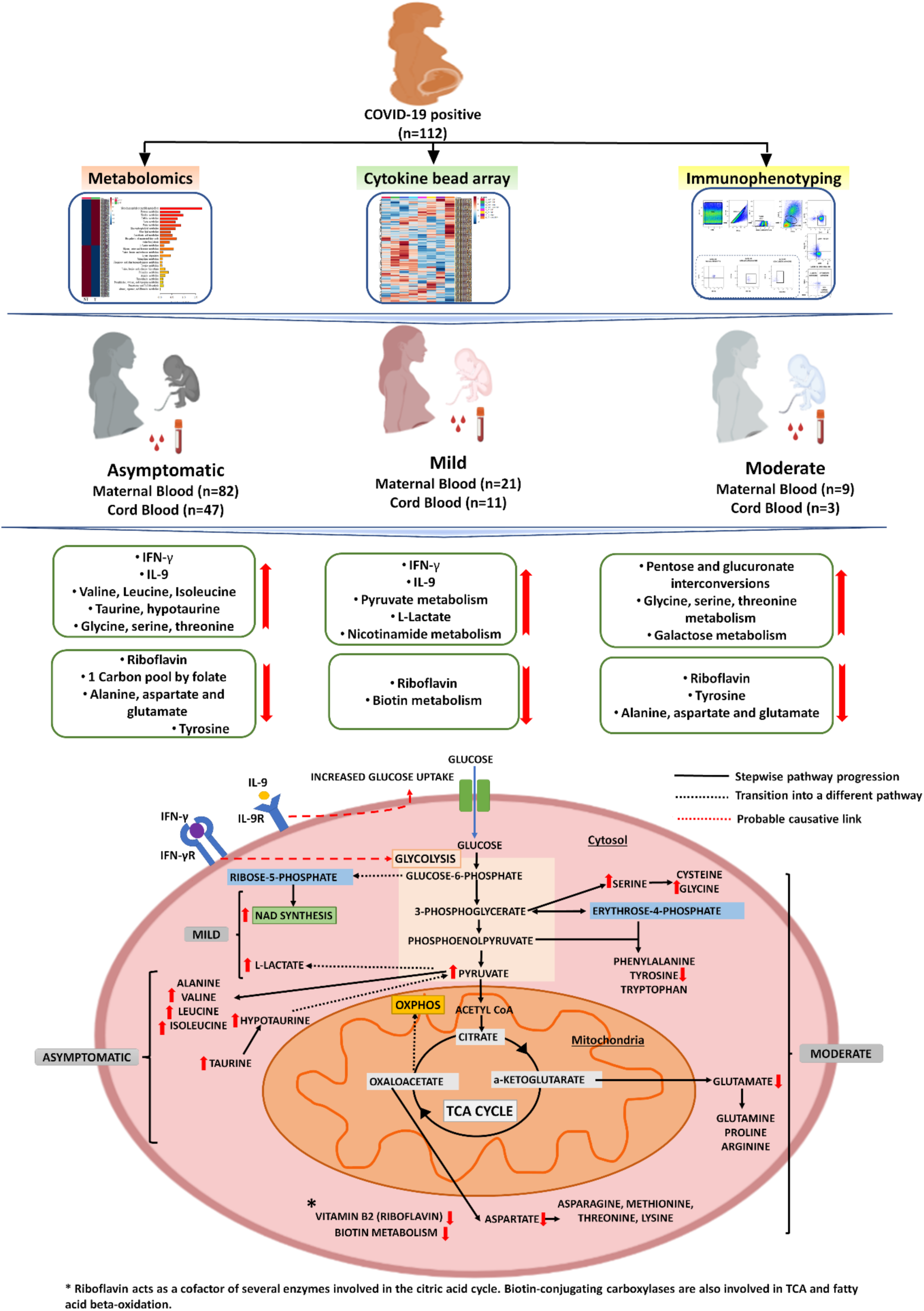

## Introduction

Many immune compromised cohorts including pregnant women are at high risk to acquire SARS-CoV-2 infection [1]. Due to vulnerability for respiratory infections [2–3] and extensively present angiotensin-converting enzyme 2 (ACE2) in placenta [4–7], pregnant women and their fetuses are more susceptible to SARS-CoV-2 infection [1]. A shift from Th1 to Th2 predominantly protects and helps the fetus and placental development but also make mothers vulnerable towards viral infections [8]. In fact, SARS-CoV-2 as highly pathogenic virus being vertically transmitted from mother to baby increase adverse pregnancy outcomes, including premature delivery, miscarriage, immune compromised neonates and maternal death [9–13].

During pregnancy, immunological state is compromised with hormonal and cytokine alterations may influence severity from asymptomatic to mild (fever, cough, mild pneumonia or absence of pneumonia) to moderate which includes hypoxia, dyspnoea, severe pneumonia which eventually leads to critical condition such as systemic shock, tachypnoea (respiratory rate, ≥30 breaths per minute) and multiple organ failure [14–15]. First trimester of pregnancy is marked with Th1- immunity characterized by production of pro-inflammatory and cell-mediated inflammatory responses, whereas in the second and third trimesters, Th9 and Tregs are involved in regulating pro-inflammatory immune responses [16–18]. Disbalanced immune responses may potentiate foetal deterioration. Furthermore, activation of T cells and extreme elimination of regulatory T cells promote uncontrolled pathogenesis in COVID-19 [19–22].

Additionally, in many viral infections like yellow fever virus, human immunodeficiency virus (HIV) [23], thrombocytopenia syndrome virus (SFTSV) [24], metabolic regulations have influence on host immunity. Similarly, in SARS-CoV-2 infection, cytokines secretion is tightly correlated with circulating metabolites [25] and elevated levels of glucose requirement was correlated with SARS-CoV-2 replication as well as enhanced cytokine production [26]. Though cytokine storms with raised IL-6, IL-1β and TNF-α have gained attention in COVID-19, but understanding of metabolic pathways regulating immune response in COVID-19 positive pregnant women is still limited. Pregnancy being immune compromised state, metabolic alteration may pose challenge to effective immune responses. Hence, these unique challenges encouraged us to investigate metabolic alterations, cytokine and immune response in SARS-CoV-2 infected pregnancy.

Herein, our study provides significant novel insights into the complex interplay between immune and metabolic responses in pregnant women with varying severity of SARS-CoV-2 infection. Our findings suggest that targeting specific metabolic pathways may present a promising avenue to address the health challenges associated with COVID-19 during pregnancy.

## Materials & Methods

### Patient recruitment

A total of 1167 pregnant women with SARS-CoV-2 infection were admitted, 1-2 days before delivery during the first and second wave of SARS-CoV-2 in Lok Nayak Jai Prakash Hospital, Maulana Azad Medical College, New Delhi. SARS-CoV-2 positivity was confirmed by nasopharyngeal RT-PCR. Before delivery, after taking informed consent, one hundred and twelve (n=112) pregnant women were recruited in our prospective study for immune-metabolomic analysis. At delivery, we were able to take cord blood samples from 82 neonates. Women with a history of COVID-19 vaccination, any comorbidity such as coronary artery disease, cardiopulmonary disease, COPD, chronic kidney disease, viral hepatitis (HBV, HCV and HEV) were excluded from this study. On the basis of clinical symptoms and disease severity, SARS-CoV-2 positive pregnant women were grouped into asymptomatic, mild, and moderate categories. Additionally, age matched healthy pregnant women (n=10) with negative SARS-CoV- 2 RT-PCR were enrolled as controls.

Out of 1167 deliveries, 33 neonates were found to be RT-PCR positive for SARS-CoV- 2 after delivery. However, from our recruited patient cohort, only 3 neonates were found to be positive for SARS-CoV-2 immediately after delivery. The study was approved by the ethics and review committee of the Institute of Liver and Biliary Sciences (ILBS) No. F.37/(1)/9/ILBS/DOA/2020/20217/585) and MAMC (No. F.1/IEC/MAMC/79/07/2020/No.186). The work was conducted in accordance with the declaration of Helsinki. Detailed methods written in Supplementary Data.

## Results

### Clinical Outcomes in COVID-19 positive pregnant women

Demographical, clinical and disease severity of 112 COVID-19 positive and healthy pregnant women is described in Table 1 Based on disease severity, out of 112 women, 82 were asymptomatic, 21 had mild symptoms (fever and cough), and 9 were having severe COVID symptoms (requiring oxygen supplementation with high fever and cough) (Supplementary Table 2). The mean maternal age was 26±5 years with median parity of one and enrolled with gestation time of 38 wks in both groups. Majority of women delivered vaginally in the healthy, whereas 62% of COVID-19 women underwent caesarean section. Apgar score was 9 at 5 minutes in the control group but only 50 babies in SARS-CoV-2 group had Apgar less than 9.

**Table 1.** Demographic and Clinical characteristics of Pregnant subjects.

|  | Healthy Pregnant<br>women as controls<br>(n=10) | SARS-CoV-2<br>positive pregnant<br>women<br>( n=112) | p Value |
| --- | --- | --- | --- |
| Age (years; median) | 25 (20-33) | 26 (21-35) | 0.85 |
| Gestational age (weeks; median) | 39(37-40) | 38 (35-40) |  |
| Parity (median) | 2 (1-3) | 1 (1-3) | - |
| Normal vaginal delivery (%) | 70% (7/10) | 38% (31/82) | - |
| Caesarean section (%) | 30% (3/10) | 62% (51/82) | - |
| Apgar score at 5 min (median) | 80% (9,9,9) | 50% (9,9,9) | - |
| Hb (mg/dL) | 11.2 (8.1-13.2) | 10.8 (6.7-14.4) | 0.32 |
| Platelet count (Lakhs) | 2.64 (1.10-2.8) | 1.63 (0.28-5.7) | 0.01 |
| TLC (μL) | 9,254(7000-13000) | 9215 (12.9-21200) | 0.1 |
| Total Bilirubin (mg/dL) | 0.59 (0.38-0.74) | 0.6 (0.1-2.8) | 0.02 |
| AST (μ/L) | 36.7 (25- 43.5) | 32.4 (10-337) | <0.001 |
| ALT (μ/L) | 38.78 (12.6-65) | 43.5 (10-390) | <0.001 |
| ALP (μ/L) | 156.5(110-200) | 222 (29-689) | <0.001 |
| SPO2 (%) | 98 (95-100) | 98 (88- 100) | 0.62 |
| D-Dimer (ng/mL) | 254 ( 125-450 ) | 560 (125-5000) | <0.001 |
| PT (sec) | 17(15-20) | 11.1 (8.6-72) | <0.001 |
| INR (sec) | 0.9 (0.8-5.2) | 0.9 (0.7-4.8) | 0.14 |
| Urea (mg/dL) | 14(12-18) | 17 (10-37) | <0.001 |
| Creatinine (mg/dL) | 0.6(0.4-0.8) | 0.5 (0.3-11.8) | 0.21 |
| Na (mmol/L) | 137(133-141) | 137 (128-147) | 0.97 |
| K (mmol/L) | 4.1(3.5-4.3) | 4.2 (3.3-7.5) | 0.01 |

COVID-19 females had significantly deranged liver function parameters with more than 1000ng/ml D-Dimer (Table 1). Interestingly, three women delivered SARS-CoV-2 positive babies, of those 2 underwent caesarean and 1 had normal vaginal delivery.

IgG concentrations of SARS-CoV-2 were determined in maternal and cord blood and found that both mothers and neonates had significantly higher levels of IgG titers than healthy (Supplementary Fig. 1).

### COVID-19 disease influences soluble factors by altering Plasma metabolomic profiles

To determine the metabolic perturbations associated with COVID-19 infection, we profiled the plasma samples using untargeted metabolomics and multivariate analysis. A total of 9,280 metabolites were extracted from raw data acquired in both positive and negative ionization modes. But after data processing and annotation, 177 metabolites were considered as differentially expressed metabolites in both groups (DEMs; *p* <0.05, FC >1.5). Volcano plot revealed that 126 metabolites were significantly downregulated and 22 were upregulated (Fig. 1A). Principal component analysis (PCA) along with hierarchical clustering was able to clearly separate the COVID-19 pregnant from healthy (Fig. 1B-C), which suggest that the metabolome profile of COVID-19 pregnant women differs from healthy pregnant women. Additionally, PLSDA (partial least square discriminant analysis) has provided few most important metabolites with VIP score (VIP; variable importance in projection) of more than 2 (Fig. 1D). Further, KEGG pathway enrichment analysis revealed that COVID- 19 infection, downregulated the tyrosine, phenylalanine and tryptophan pathways associated metabolites (Fig. 1E) but upregulated the amino acids such as valine, leucine and isoleucine and metabolites associated with glycine, serine, threonine and pyruvate metabolism (Fig. 1F). Our results highlight that COVID-19 disease most strongly affects tyrosine and tryptophan associated energy pathways.

**Figure 1.**
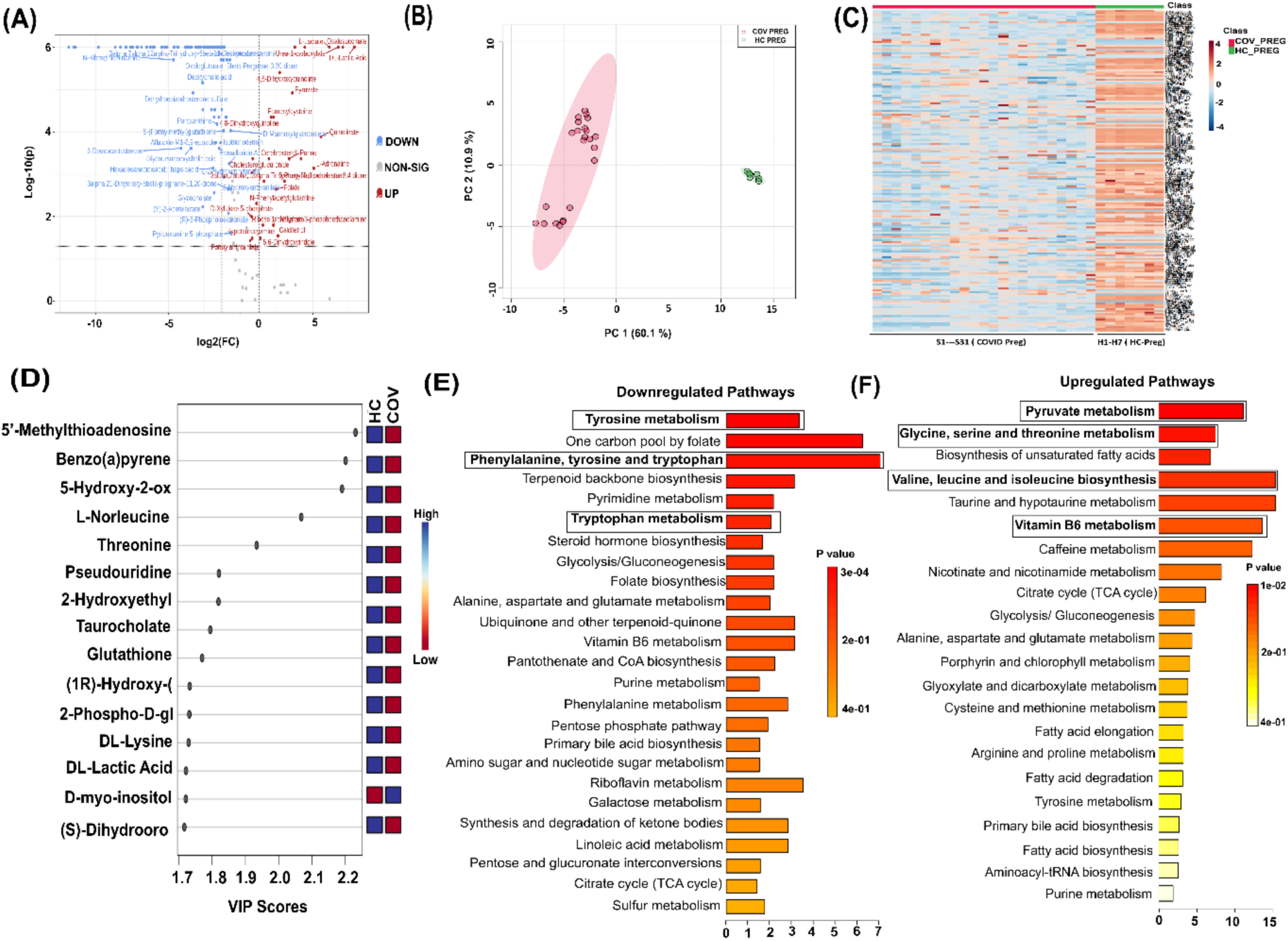
Plasma metabolomics profile of COVID-19 positive and healthy pregnant women. **(A)** Volcano plot showing differentially expressed metabolites in two groups. Red dots represent upregulated, blue dots represent downregulated (p < 0.05) and gray dots represent non-significant metabolites **(B)** Partial component analysis (PCA) showing clear segregation between two groups. Green dots correspond to healthy control and pink dots correspond to COVID-19 positive pregnant women **(C)** Heat map analysis are capable to segregate COVID-19 positive pregnant women (red bar) from healthy control pregnant women (green bar) The expression is given as red upregulated and green downregulated **(D)** Variable importance in projection (VIP) plot displaying the top 15 most important metabolite features identified by PLSDA. Colored boxes on the right indicate relative concentration of the corresponding metabolite between two groups. **(E-F)** upregulated and downregulated metabolites associated with KEGG enriched pathways.

### Decrease in amino acid metabolism but increase in nicotinamide- nicotinate metabolism is associated with disease severity

To delineate the change in metabolome with increased disease severity, we have compared SARS-CoV-2 positive pregnant women with different severity and with healthy. Volcano plots have provided the differentially expressed metabolites in each subgroup comparison and PLSDA has provided a list of most important metabolite features among all subgroups (Supplementary Fig. 2). Further, pathway analysis of metabolite profiles of subgroups revealed significantly enriched pathways (FDR<0.05). In asymptomatic SARS-CoV-2 positive pregnant, valine, leucine, isoleucine, taurine, hypotaurine, glycine, serine and threonine metabolic pathways. were altered compared to healthy. In this group, valine, leucine, isoleucine and taurine, hypotaurine metabolism, glycine, serine and threonine metabolism pathways were significantly upregulated and riboflavin, alanine, aspartate and glutamate and tyrosine pathways were significantly downregulated (Fig. 2A-B). Riboflavin was commonly found downregulated in the healthy vs. asymptomatic group and mild to moderate group. Whereas, with increase in disease severity (mild group), nicotinate and nicotinamide metabolism was upregulated but pyruvate and biotin metabolism was downregulated (Fig. 2C-D).

**Figure 2.**
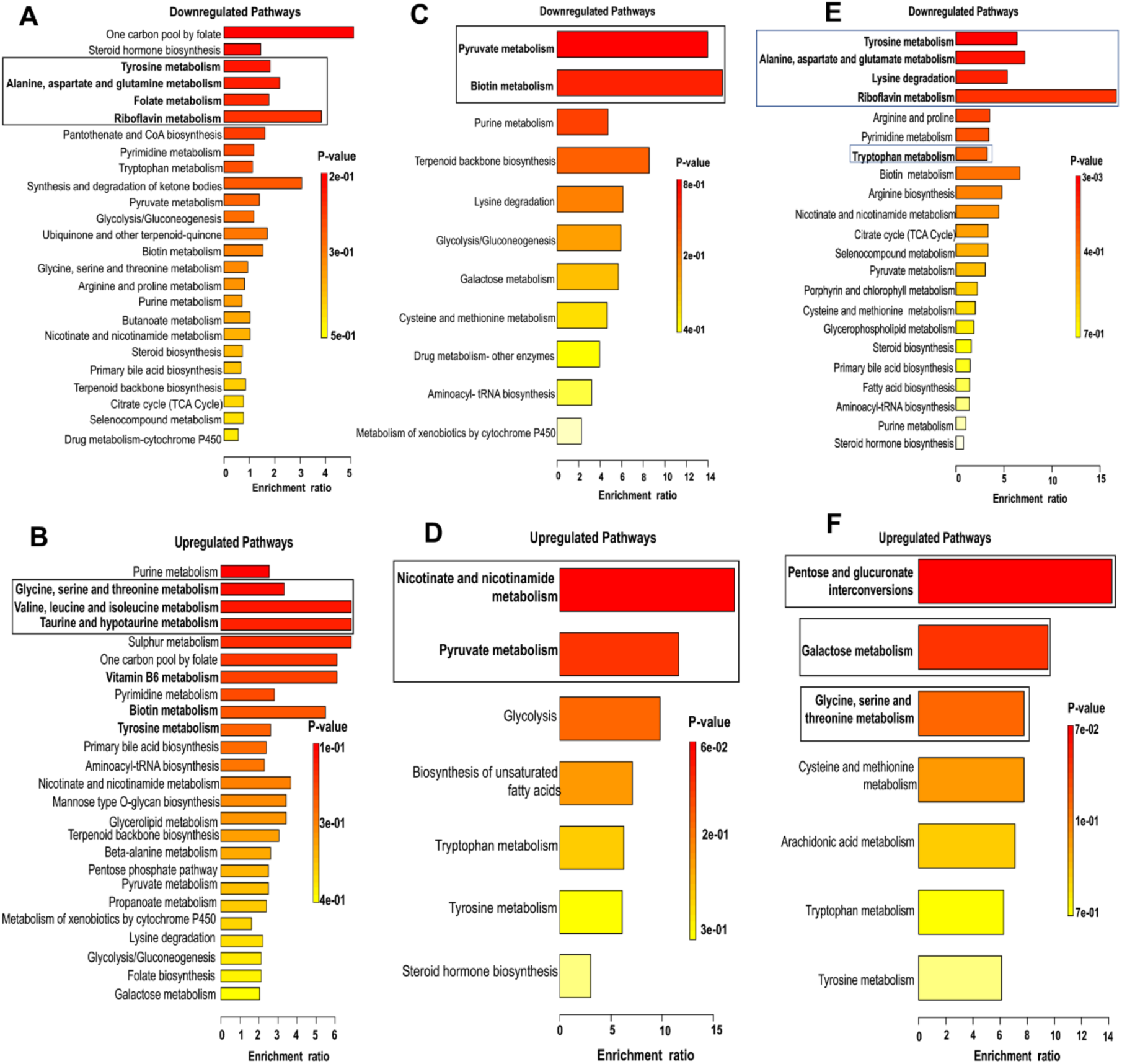
Pathway enrichment using KEGG in SARS-CoV-2 positive pregnant women with disease severity. KEGG enrichment reveals downregulated and upregulated pathways; **(A-B)** Asymptomatic pregnant women compared to healthy **(C-D)** Pregnant women with mild disease compared to Asymptomatic **(E-F)** Pregnant women with moderate disease compared to mild.

Furthermore, from mild to moderate shift (Fig. 2E-F), riboflavin, tyrosine and alanine, aspartate and glutamate pathways were downregulated but upregulation in glycine, serine, threonine, galactose metabolism and pentose with glucuronate interconversions were observed (Supplementary Table 3).

### Resemblance of metabolic signatures in mothers and their neonates

To understand the metabolic changes in neonates born from mothers with disease severity, we compared the metabolic profiles of umbilical cord bloods, as the cord blood provides a representative of a newborn’s blood. Pathway analysis of metabolite profiles of subgroups revealed significantly enriched pathways (FDR<0.05).

Similar to maternal blood, cord blood from asymptomatic pregnant females showed upregulation of valine, leucine, isoleucine, glycine, serine and threonine metabolic pathways, and nicotinamide and nicotinate metabolism, accompanying downregulation of alanine, aspartate and glutamate and tyrosine pathways (Fig. 3A- B).

**Figure 3.**
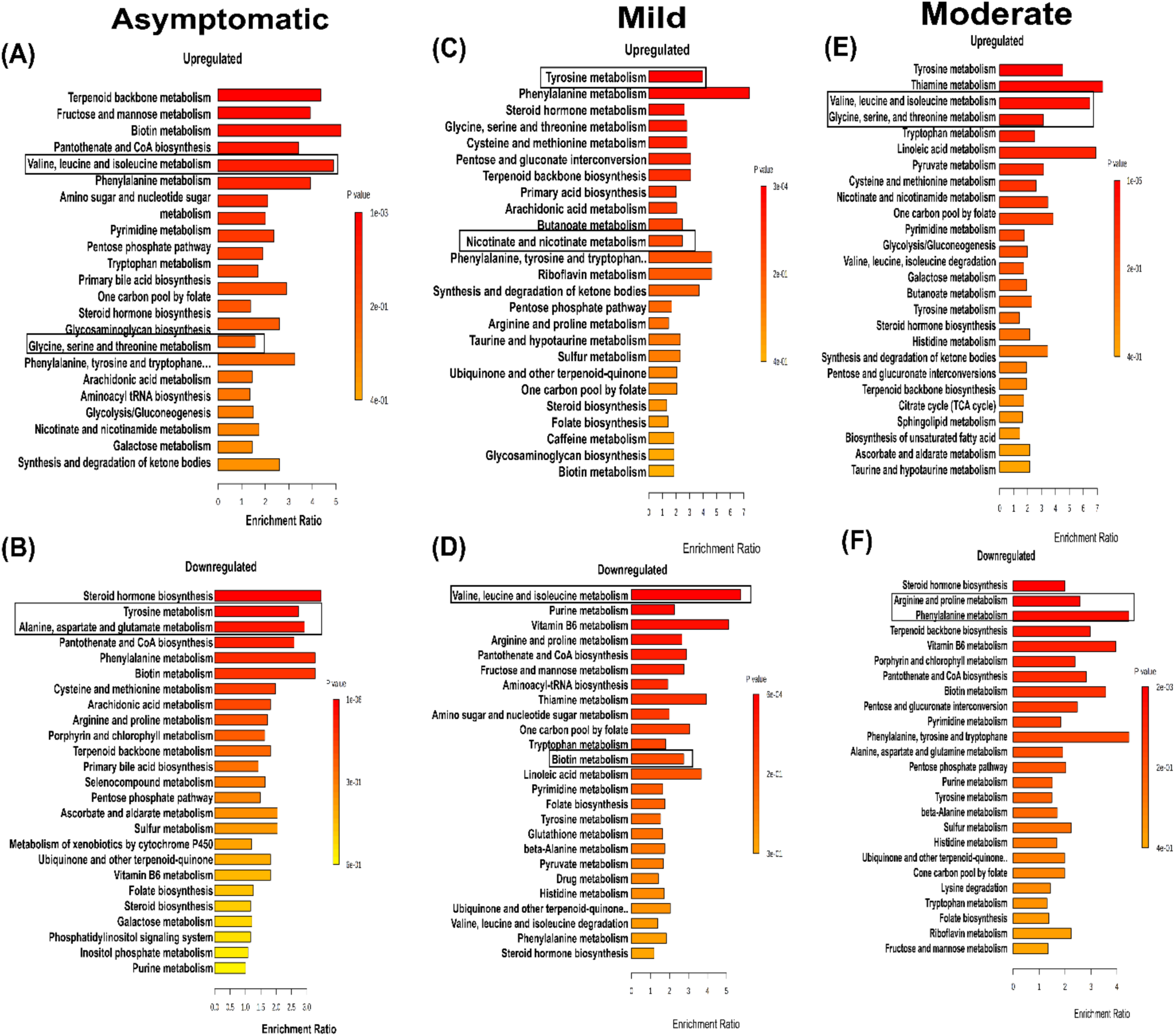
Pathway enrichment using KEGG in cord blood from SARS-CoV-2 infected pregnant women with disease severity. KEGG pathway enrichment analysis reveals upregulated and downregulated metabolic pathways; **(A-B)** Asymptomatic pregnant women **(C-D)** Pregnant women with mild disease **(E-F)** Pregnant women with moderate disease.

Whereas metabolites in cord blood from mild group had upregulation of tyrosine and nicotinamide and nicotinate metabolism and valine, leucine, isoleucine and biotin metabolism were downregulated (Fig. 3C-D).

As, severity increased in moderate group amino acid metabolism was upregulated including tyrosine, valine, leucine, isoleucine, glycine, serine, threonine and pentose and gluconate interconversion metabolic pathways. However, arginine, proline, phenylalanine and riboflavin metabolism were downregulated (Fig. 3E-F).

Our results suggests that cord metabolic pathways were almost mimicking their mothers’ profile with little variation.

### COVID-19 infection causes cytokine storm in mothers, their babies’ cord and placenta

The cytokine storm (a severe immune response) has been recognized as a major contributor to multi-organ failure in COVID-19 patients. Recent studies revealed cytokine storms in *SARS-CoV-2 positive* pregnant women [28–29]. However, our results from both mother and cord duo blood samples revealed significantly raised levels of pro-inflammatory cytokines. Our nineteen cytokine panel (Supplementary Table 1) showed significantly increased concentrations of TNF-α, IFN-α, IFN-β, IFN- γ, IL-1β, IL-17a, IL-21, IL-22, IL-29, and IL-33 (pro-inflammatory) IL-2, IL-4, IL-7, IL-9, and IL-15 (anti-inflammatory) cytokines in COVID-19 infected pregnant (Fig. 4A). Results from twenty three cord blood samples (15 from COVID-19 infected mothers and 8 from healthy) reveled the raised levels of pro and anti-inflammatory cytokines including TNF-α, IFN-α, IFN-γ, IL-17a, IL-21, IL-22, IL-29, IL-33, IL-4, IL-6, IL-9, IL-10, and IL-15 in infected samples than healthy (Fig. 4B). Basically, except IL-1β, all other cytokines representing cytokine storm including TNF-α, IFN-α, IFN-γ, IL-6, were significantly elevated in both COVID-19 mothers and cord blood.

**Figure 4:**
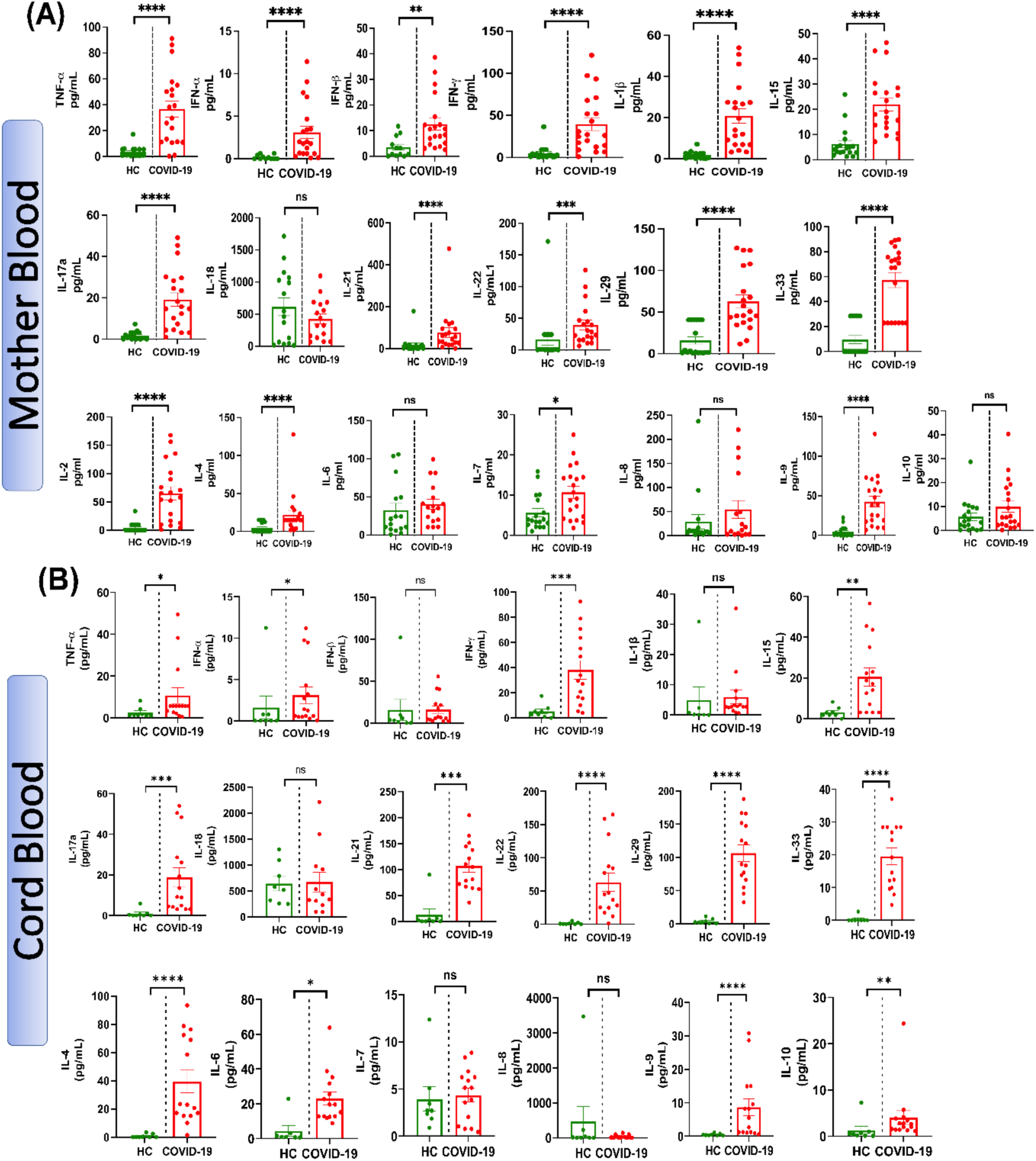
Cytokines concentration in COVID-19 positive mothers and in cord blood of their neonates. **(A)** Plasma concentration of TNF-α, IFN-α, IFN-β, IFN-γ, IL- 1β, IL-17a, IL-18, IL-21, IL-22, IL-29, IL-33, IL-2, IL-4, IL-6, IL-7, IL-8, IL-9, IL-10, and IL-15 from the peripheral blood of SARS-CoV-2 positive (n=20) and healthy control (n=18) women. **(B)** Plasma concentration of TNF-α, IFN-α, IFN-β, IFN-γ, IL-1β, IL-17a, IL-18, IL-21, IL-22, IL-29, IL-33, IL-2, IL-4, IL-6, IL-7, IL-8, IL-9, IL-10, and IL-15 from the cord blood of SARS-CoV-2 positive (n=15) and healthy control (n=8) women. Data are shown as boxplots where red dots indicate SARS-CoV-2 positive (Covid-19) women. The data are presented as mean ± standard deviation (SD) and analyzed using unpaired t-test with Welch’s correction. Significance levels are indicated as *p<0.05, **p<0.01, ***p<0.001, and ****p<0.001.

Additionally, pro- and anti-inflammatory cytokines levels were analysed with disease severity; asymptomatic (n=6), mild (n=7), and moderate (n=7) subgroups of SARS- CoV-2 infected pregnant women. Though the women were asymptomatic, but with COVID-19 infection, levels of most of the pro-inflammatory cytokines including TNF- α, IFN-β, IFN-γ, IL-1β, IL-17a, IL-21, IL-22, IL-29, IL-33 and anti-inflammatory cytokines IL-2, IL-9, and IL-15 were similarly elevated as in mild (Supplementary Fig. 3). With moderate disease, there is more immunosuppression with increased IL-4 levels. Notably, IFN-α and IL-10 were decreased in the moderate group compared to healthy and mild. When we compared cord blood profiles in asymptomatic (n=5), mild (n=6), and moderate (n=4), it was observed that IL-22, IL-29, and IL-8 were significantly increased in both the asymptomatic and mild cords. Besides that, IL-9 was the only cytokine found to be significantly elevated in all three groups (asymptomatic, mild, and moderate) compared to healthy. With mild disease, most of the cytokines including TNF-α, IFN-α, IFN-β, IFN-γ, IL-1β, IL-17a, IL-21, and IL-33 (pro-inflammatory cytokines), and IL-2, IL-6, IL-7, IL-10, and IL-15 (anti-inflammatory cytokines), were significantly elevated in the cords (Supplementary Fig. 4). Further, we have also analysed TNF-α, IFN-αR in placental tissue using qRT-PCR and observed that TNF-α expression was fivefold increase in COVID-19 infected placenta than healthy (Supplementary Fig. 5). IFN-αR was also increased but difference was not significant. Therefore, our result suggests that SARS-CoV-2 infection in mothers pose deleterious effects on the fetus in the womb.

### COVID-19 infection alters the immune signatures by increasing *expression of* HLA-DR, CCR2 and CX3CR1 on *monocytes*

Whole blood immunophenotyping revealed that there was no change in total monocyte, classical and intermediate monocytes except increase in non-classical monocytes in COVID-19 (Fig. 5B). There was also no change in mean fluorescence intensity (MFI) of CD16 or CD14 (data not shown) compared to healthy. Considering classical monocytes with very short circulating lifespan, viral proteins containing classical monocytes may turn into intermediate or non-classical monocytes, which indeed was increased in patients compared to healthy (Fig. 5B). Although, with disease severity, monocyte population was decreased in pregnant woman with moderate disease but difference was not significant may be due to less patient numbers in this group (Fig. 5C). Further, HLA-DR, CCR2 and CX3CR1 expression was definitely increased in classical monocytes indicating their migration and phagocytic ability in circulation (Fig.5D). We did not find any significant HLA-DR, CCR2 and CX3CR1 expression in non-classical and intermediate monocytes. These results signify that non-classical monocytes are majorly involved in providing first line defense in Covid-19 infected pregnant females. Other than this, pregnant women with SARS-CoV-2 infection show a considerable decreased numbers of conventional DCs with increase in myeloid DCs (Supplementary Fig. 6).

**Figure 5:**
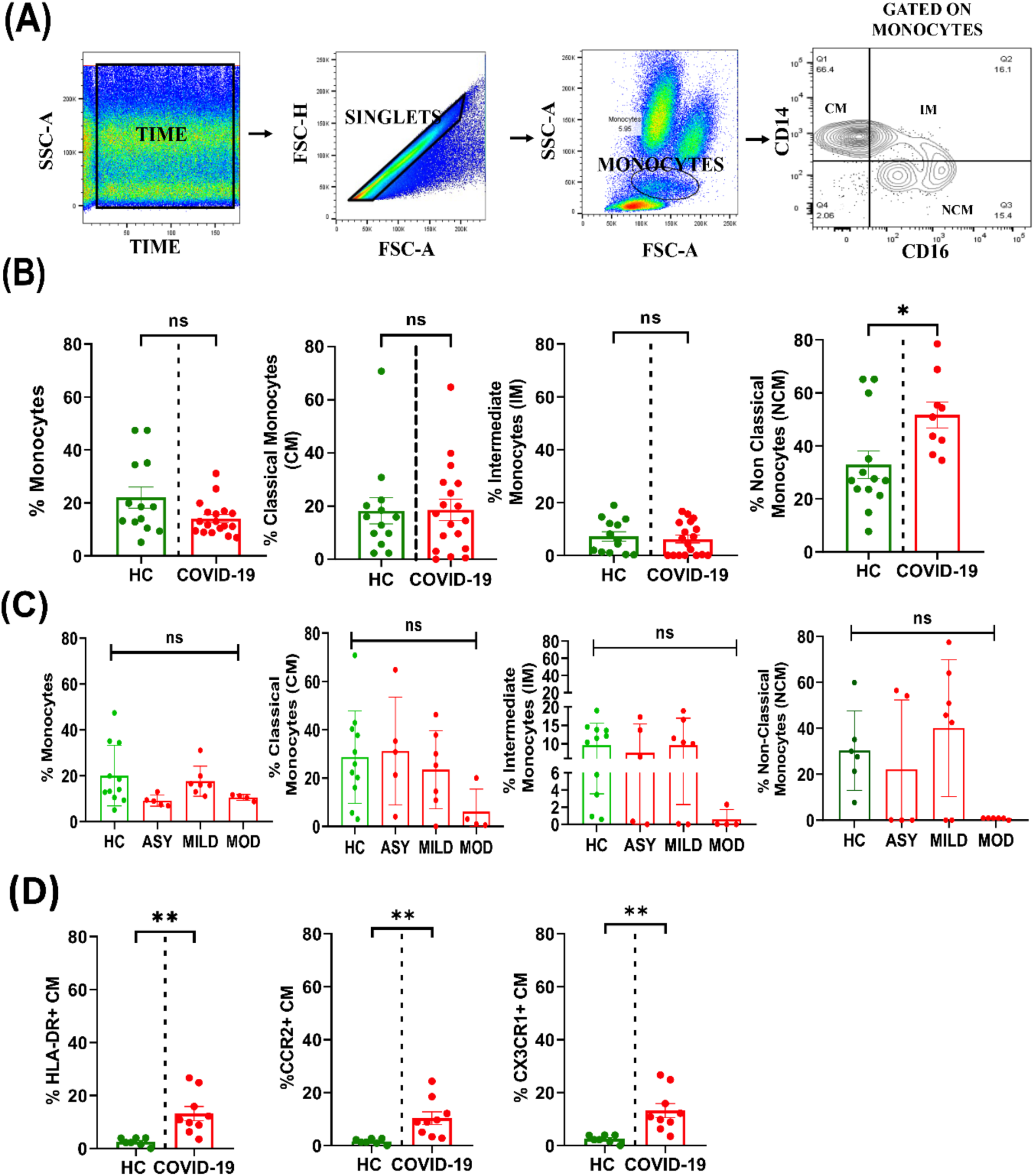
Total monocytes and their subset population in COVID-19 pregnant samples. **(A)** Gating strategy of monocytes and monocyte subsets on the basis of CD14 and CD16 positivity (Classical monocytes (CM) CD14+CD16-; Intermediate monocytes (IM) CD14+CD16+ and non-classical monocytes (NM) CD14-CD16+) **(B- C)** Percentage frequency of total monocytes and subsets in COVID-19 patients and its subgroups (asymptomatic, mild, and moderate) **(D)** Percentage expression of HLA-DR, CCR2 and CX3CR1 on classical monocytes in COVID-19 positive pregnant women. Data is presented as mean ± standard deviation (SD) and analyzed using unpaired t-test with Mann-Whitney correction and one way ANOVA. Significance levels are indicated as *p<0.05 and **p<0.01.

### Decreased memory B cell frequencies in COVID-19 infected pregnant women

Our results revealed that there is no significant difference in total B cell frequency in COVID-19 positive population than healthy (Fig. 6B). But in subgroups (asymptomatic, mild, and moderate) this difference was observed especially with moderate disease (Fig. 6B). Further, naïve, immature naive B cells were increased and memory B cells were decreased in COVID-19 pregnant (Fig. 6C). However, with few memory B cells, still COVID-19 pregnant struggled to have reasonable numbers of switched memory B cells and plasma blasts than healthy (Fig.6D). In COVID-19, B cells gained significant expression of CD40 and somewhat higher expression of A proliferation inducing ligand (APRIL) and B-cell activating factor receptor (BAFF-R) than healthy (Fig. 6E).

**Figure 6.**
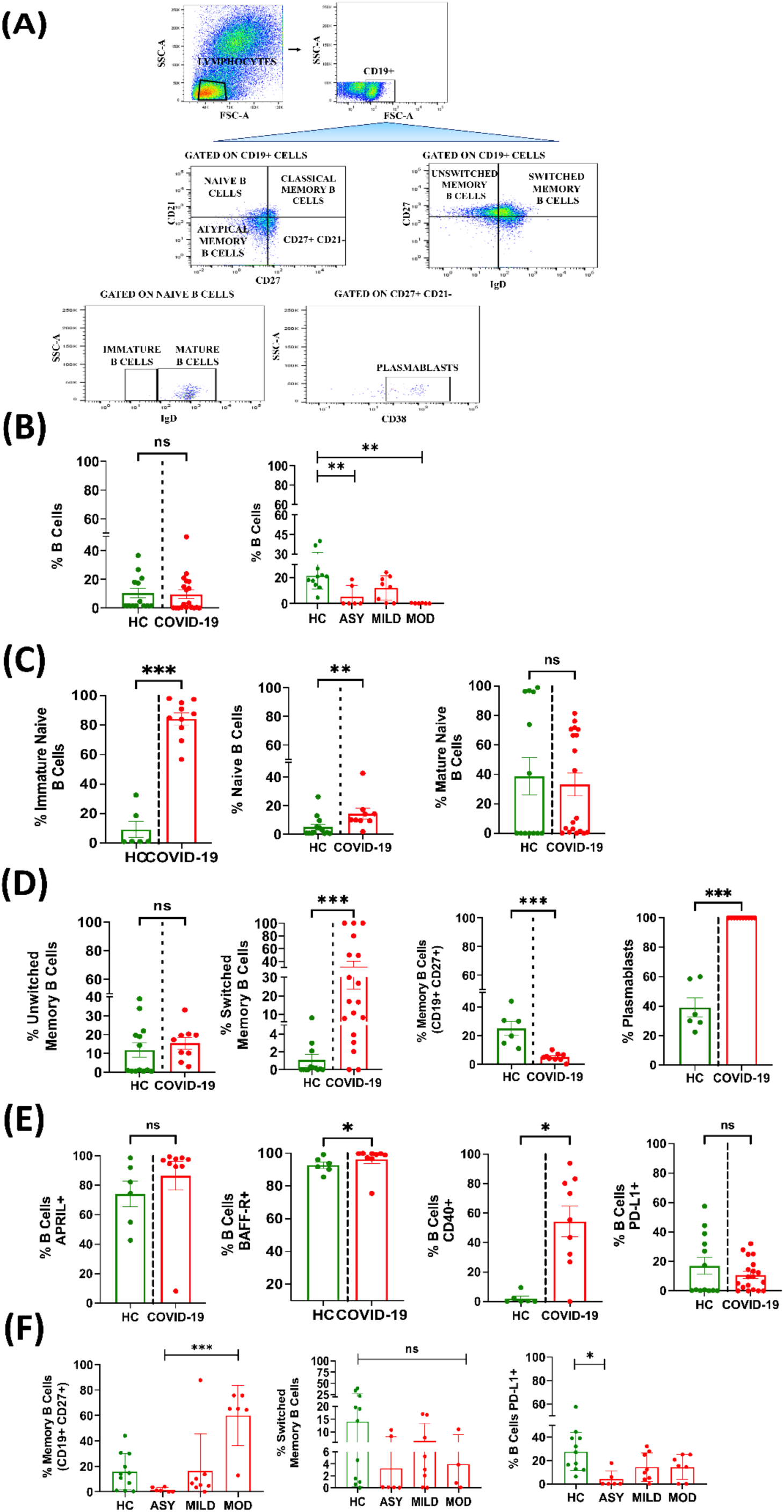
B cells compartment in COVID-19 pregnant patients. **(A)** Gating strategy for B cells and their subsets **(B)** Percentage frequency of total B cells in COVID- 19 and their subgroups **(C)** Immature naive, naive and mature naive B cells **(D)** Un- switched memory, switched memory, memory B cells and plasmablasts **(E)** Expression of APRIL, CD40 and PDL-1 on total B cells **(F)** Frequency of memory B cells, switched memory B cells, and PDL-1+ B cells in COVID-19 positive pregnant women with disease severity. Data is presented as mean ± standard deviation (SD) and analyzed using unpaired t-test with Mann-Whitney correction and one way ANOVA. Significance levels are indicated as *p<0.05, **p<0.01, and ***p<0.001.

Further, we observed that exhaustive cell surface markers on B cells and observed that in COVID-19, B cells had no differential expression of PDL1, however when compared in subgroups, it was noticed that asymptomatic group had less exhaustive B cells than mild and moderate (Fig. 6F). Along with PDL1, together our results highlight that indeed B cell compartment is compromised in COVID-19 positive pregnant women which may have implications in developing fetus long lasting immunity.

### Raised CD4+T cells contribute to existing immune compromised state of pregnancy

COVID-19 disease has affected T cells in pregnant women especially CD4+T helper cells (Fig 7B). We have observed a significant elevation in CD4+ T naïve cells (CD4+ CD45RA+ CCR7+), TCM (central memory), TEM (effector memory) in the disease condition (Fig. 7C), in contrast, there was no significant change in cytotoxic CD8+T cells and specifically had low numbers of naïve, central memory, effector memory CD8 T cells but with increased frequencies of CD8 TEMRAs (terminally differentiated effector memory) (Fig. 7D). With disease severity, even CD4 T cells were affected and in moderate condition CD4 naïve and CD8 EM were decreased than mild, although these differences were not significant may be due to less patients in moderate group (Fig. 7E). Pregnancy is more of with Th2 state, but our results suggest, SARS-CoV-2 infection further raise CD4^+^ T helper cells and add into immune suppressed state.

**Figure 7:**
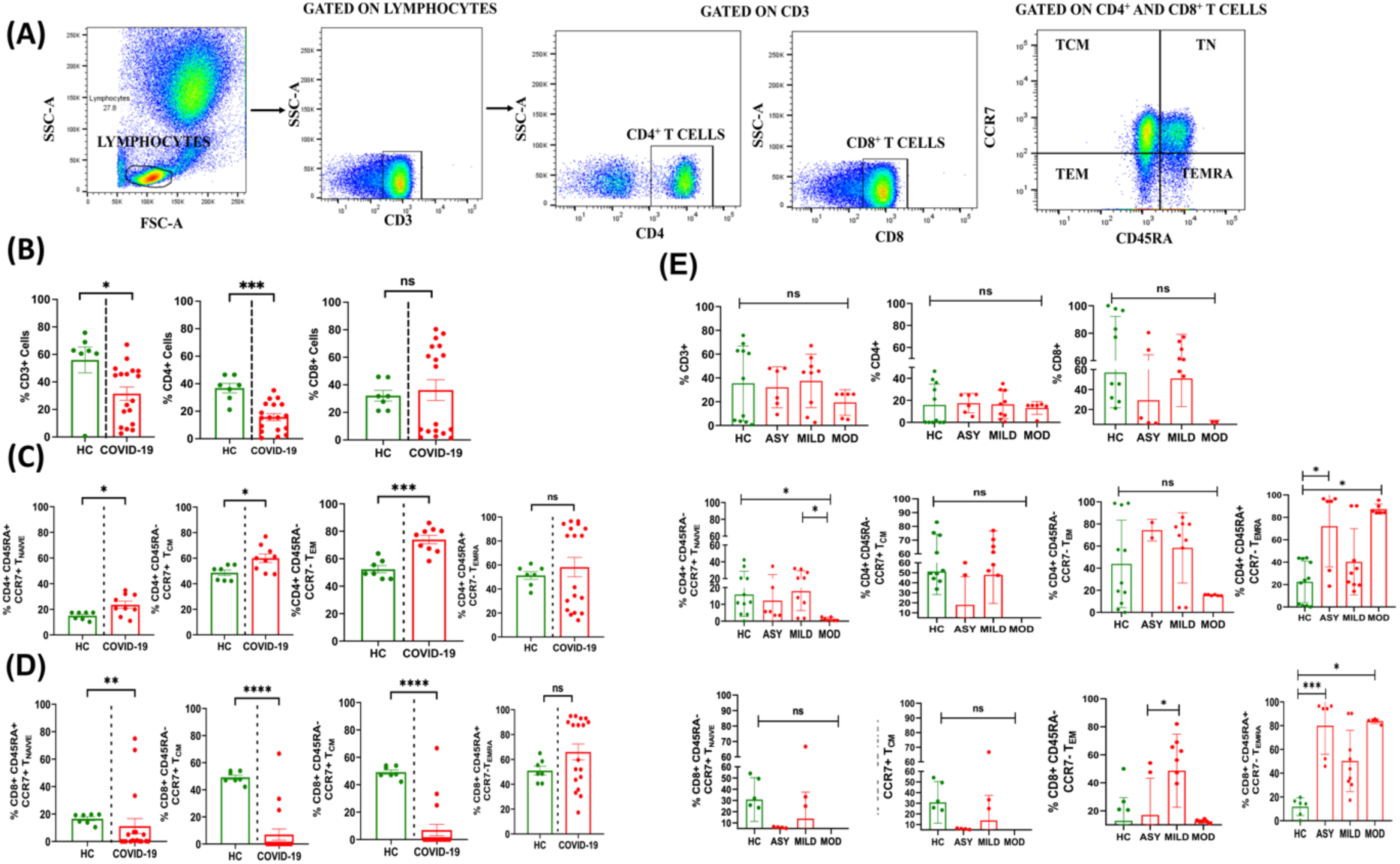
T cells and their subsets in COVID-19 pregnant patients and healthy control. **(A)** T cell gating strategy **(B)** Percentage frequencies of total CD3+, CD3+CD4+ and CD3+CD8+ T cells **(C)** CD4+ Tcells along with their naïve, central memory, effector memory and TEMRA subsets. **(D)** CD8+ cells along with their naïve, central memory, effector memory and TEMRA subsets. **(E)** T cells in Asymptomatic, mild and moderate pregnant patients. The data are presented as mean ± standard deviation (SD) and analyzed using unpaired t-test with Mann-Whitney correction and one way ANOVA. Significance levels are indicated as *p<0.05, **p<0.01, ***p<0.001, and ****p<0.0001.

Our analysis revealed that the CD8^+^ T cell population in SARS-CoV-2 infected pregnant women was exhausted, as evidenced by decreased levels of CD8^+^ naïve T cells, TCM and TEM cells.

## Discussion

COVID-19 pandemic continues to persist with the emergence of new variants and pose threat to immune compromised cohorts including pregnant women. Therefore, it was crucial to understand the alteration in cytokine profiles, metabolomic landscape influencing immune responses in pregnant mothers and their long-lasting effects on their fetus.

Our data revealed that along with raised IL-6, TNF-α, IL-1β cytokine storm, which impacts on organ and disease pathogenesis [27–28], COVID-19 disease also increased IFN-γ, IL-9, L-21, IL-22 and IL-33 in all pregnant women irrespective of their disease severity. Generally, in pregnancy IL-9 levels are increased however, it is slowly decreased till the time of delivery [29], however though we have observed raised IL-9 levels in COVID-19 infected women at the time of delivery. Since IL-9 participates in host-pathogen interaction and immune mediated disease [30–31], its increased levels may pose potential threat to the fetus.

Plasma metabolic analysis revealed that along with increased IL-9 and IFN-γ, most of the metabolites associated to energy consumption including tyrosine, valine, leucine, isoleucine, L-lactate, amino acid metabolism, pyruvate metabolism, and riboflavin was were altered in COVID-19 pregnant. Interestingly, we observed increased levels of TNF-α and IL-9 in diseased mothers’ cord blood and also increased expression of TNF- α in placental tissue. Indeed, we have observed similar metabolites in COVID-19 exposed cord blood samples. However, out of 112 deliveries, we found only three babies with SARS-CoV-2 infection, but metabolomic results suggests that, though not all infants were infected but in utero exposure to SARS-CoV-2 have altered the metabolomic profile as it did in mothers.

In asymptomatic pregnant women, with increased IL-9 and IFN- γ, valine, leucine, isoleucine, glycine, serine and threonine were upregulated which increases glucose uptake and glycolysis [26]. Further, increased levels of IL-9 produce more of L-lactate which shows its involvement in TCA cycle by elevating lactate levels [31]. Therefore, these findings suggest IL-9 as a potential biomarker for the prediction of COVID-19 severity in pregnant women during the third trimester and at the time of delivery.

When disease reaches a mild condition, the body feels lethargic and required more energy which is being compensated with consumption of stored glucose by upregulating pyruvate and nicotinamide and nicotinate metabolism. Moreover, IFN-γ found to be closely regulating cellular metabolic pathways and requires high rates of glycolysis [26, 32] which increases the production of pyruvate and NAD^+^. Therefore, it can be comprehended that IL-9 and IFN-γ both dysregulate the TCA cycle by increasing the levels of lactate and pyruvate in COVID-19 infected pregnancy. Our results revealed that cord blood metabolic pathways were mimicking with their mother’s profile. Altered metabolism in cord blood may cause adverse effects in neonates, although the long-term consequences following in utero exposure are unknown. However, neonates exposed to SARS-CoV-2 appears relatively benign, some infants might develop respiratory distress and require supplemental oxygen or intubation [33–34].

Metabolic regulations severely effects immune compartment especially in acute inflammations. Human monocytes from bone marrow come into circulation as classical monocytes but due to short life span quickly transitioned into intermediate and non-classical monocytes [35]. Classical monocytes have very short life span and especially after interacting with viral proteins, classical monocytes turn into intermediate and non-classical monocytes. It was observed previously that, increases in glycolysis and pentose phosphate pathways induce activation and transitions of monocytes [36]. Indeed, COVID-19 pregnant group showed increased non-classical monocytes despite the total decreased monocytes with disease severity. In our study, monocytes showed increased expression of CCR2 and previously, it was observed that increased concentrations of CCL2 initiated cytokine storm and promoted myeloid cells infiltration in the lungs airways which caused alveolar damage in COVID-19 patients [37]. In accordance to this study, our findings show similar results with an increase in CCR2 and CXC3R in SARS-CoV-2 infected pregnant women. Therefore, CCR2 and CXCR3 play key role in immune pathogenesis of COVID-19 in pregnant women.

With increased Il-9, IFN-α levels, we have also observed more of TNF-α in mother and cord blood as well as higher expression of TNF-α in placental tissue especially in the mild group (Supplementary fig 6). TNF-α is considered as a potent regulator of lipid and NAD^+^ metabolism [38]. Moreover, in accordance with the earlier reports, increased levels of TNF-α and Il-6 may induce preeclampsia in pregnancy [39]. Preeclampsia is also characterized by increased levels of ESR2 and GPER1 [40] and indeed we have observed elevated expression of ESR2 and GPER1 in COVID-19 infected placental tissue (Supplementary Fig. 7), however, in our cohort we have observed only a single case of preeclampsia. However, these results suggest that immune activation and exhaustion both pose threat to newborn with long-term neurodevelopmental problems.

In moderate to severe stages, downregulation of riboflavin shows more energy consumption by cells. These results signify that more energy is required in COVID-19 mild condition as NAD^+^ is used for consumption of stored food or glucose. Moreover, at severe condition of COVID-19 in pregnancy, downregulation of riboflavin causes hindrance to perform its functions to activate innate immunity, improving respiratory functions and reducing pro-inflammatory cytokine levels.

T cells play a major role in viral clearance in which CD4+ helper T cells (Th2) supports CD8+ cytotoxic T cells, which secrete granzymes, IFN-γ and perforin to eradicate viruses from the host [41]. Elevated CD4+ naive T cells in the diseased group, found to be associated with the production of IL-4 cytokine by SARS-CoV-2 infection, leading to T cell differentiation into the Th2 subset. Furthermore, NAD^+^ is characterized to control T cell survival [42–43], hence, increased levels of NAD^+^ induce apoptosis in CD8+ T cells. In our study, CD8 T cell frequencies in COVID-19 positive pregnant were decreased and TEMRA T cells were significantly elevated in asymptomatic and moderate groups. In moderate disease, exhausted T cell eventually loose production of IL-2 and IFN-γ cytokines. Taken together this concludes that immune response in pregnant women varies with disease severity. However, in the mild group compared to asymptomatic CD8 TEM were significantly increased, indicating a possible stronger immune response in mild cases. These findings suggest that disease severity can affect the T cell response to COVID-19 in pregnant women, which may have implications for the development of long-lasting immunity and the risk of reinfection.

These findings suggest that altered metabolites involved in SARS-CoV-2 infected pregnancy may alter the immune responses which may serve as drivers of COVID-19 related complications during third trimester of pregnancy or at the time of delivery. In moderate disease, vitamin B2 (riboflavin) dependency and downregulation cause more metabolic usage in B cell [44] leading to decrease in mature naive B cells and memory B cells which eventually leads to decrease in production of IgA, by which inflammatory responses and secretory immune responses also decreases.

As the B memory cells secrete antibodies and mediate the adaptive immune response making them more important to protect immunity against COVID-19 disease [45–46]. In the present study, frequency of memory B cells were decreased in COVID-19 positive pregnant group but still their maturation capacity was good to make plasmablasts, which diminishes in mild to moderate conditions. Though, deeper insight is needed for SARS-CoV-2 specific longevity and nature of B memory cell compartment but our results indicate that apart from memory B cells, naive B cells, immature B cells, central memory B cells and plasmablasts have contributed to increase humoral immune response during pregnancy with SARS-CoV-2 infection. We do have limitation in our study, as our results were based on maternal blood and cord blood duo samples from pregnant woman with varying disease severity. But due to logistic issues, cord blood samples were not processed for immune phenotyping.

## Author’s contributions

SH: Performed metabolomics experiments, data analysis, interpretation and manuscript writing PP: Performed Immune experiments acquisition and analysis, metabolomics data analysis HS: Patient material collection and initial processing, MK and AS: Recruitments of COVID-19 pregnant patients JKS RS and AS: Performed cytokine bead array MJ; RNA isolation and Quantitative RT-PCR from placentas PY: helped in drawing graphical abstract PK; IgG determination in samples SKS: Manuscript editing and intellectual inputs JSM: helped in metabolomics data analysis ST: COVID-19 patient stratification, clinical data analysis and clinical inputs in manuscript NT: Conceptual design of study, designing of all experiments, Immune phenotyping, data interpretation, critical revision of the article for important intellectual content and finalized the manuscript.

## Acknowledgments

We thank our technicians, Mr. Dileep and Mr. Abhishek Tomar at MCM for the storage and initial processing of few samples.

