## Supplementary Data for "Metabolic alterations unravel the materno–fetal immune responses with disease severity in pregnant women infected with SARS-CoV-2"

**Affliations:**

^1^Laboratory of Molecular Immunology, Department of Molecular and Cellular Medicine, Institute of Liver and Biliary Sciences, New Delhi, India

^2^ Department of Gynecology and obstetrics, Lok Nayak Jai Prakash Hospital, New Delhi, India

^3^ Department of Microbiology, Institute of Liver and Biliary Sciences, New Delhi, India

^4^ Department of Hepatology, Institute of Liver and Biliary Sciences, New Delhi, India

*Corresponding authors

**Methods:**

Clinical parameters and relevant investigations were collected from patient admission

file and hospital records. Maternal blood, cord blood and placenta were collected at

the time of delivery.

**Laboratory Investigations and Analysis**

***Blood Sampling:***

About 9-10 ml of peripheral blood and 5-6 ml of cord blood was collected in EDTA tubes. 100-120µl of whole blood was used for immune phenotyping using high dimensional flow cytometry panels and rest of the blood was spin down to collect platelet free plasma which was stored at -80^o^ C for various biochemical, serological, metabolomics analysis as well as for cytokine-bead array assay. After separating plasma, rest of the blood sample was used for peripheral blood mononuclear cells (PBMCs) isolation.

***Serological estimations:***

Complete blood count (CBC) using hematology analyzer, and transfusion-transmitted infections like Hepatitis C virus (HCVcAg), Hepatitis B virus (HBsAg), HIV, and malaria were done as per manufacturer protocol and analyzed by a fully automated ARCHITECT i2000SR immunoassay analyzer (Abbott Laboratories).

***Detection of SARS-CoV-2 specific antibodies:***

The SARS-COV-2 IgG antibodies were measured using the enhanced chemiluminescence method (Vitros ECi, Ortho Clinical Diagnostics, New Jersey, US). This a qualitative assay based on a recombinant form of the spike subunit 1 protein. Results are based on the sample signal-to-cut-off (S/Co) ratio, with values <1.0 as negative and ≥1.00 as positive results.

***Analysis of plasma metabolites:***

*Sample preparation:*

We have analysed plasma metabolites in healthy pregnant women (n=10) and COVID-19 pregnant women (n=56) their cord bloods (n=24). In brief, 100μl of plasma samples was used for metabolites extraction using organic phase extraction method. In brief, 400 μl methanol was added to 100 μl plasma, and then vortexed/mixed properly and kept overnight at −20°C for [protein precipitation](https://www.sciencedirect.com/topics/chemistry/protein-precipitation). The next day, samples were centrifuged at 13000 rpm for 10 minutes at room temperature. Supernatant was collected and dried under vacuum.

*Mass Spectrometric Analysis of metabolites mixtures*

The dried samples were reconstituted with 95% water, 5% acetonitrile and 5% internal standards (In house lab protocol) in the ratio of (95:5:5) for reverse-phase chromatography. C18 column (Thermo Scientific 25003102130: 3 μm, 2.1 mm, 100 mm) was used for ultra-high performance liquid chromatographic (HPLC) system followed by high-resolution mass [spectrometry](https://www.sciencedirect.com/topics/engineering/spectrometry) (MS).

*Data processing:*

Compound discoverer 3.0 software was used to extract metabolite features followed by performing annotation of features using mzCloud ([www.mzcloud.org](http://www.mzcloud.org/)) and [mummichog](https://www.sciencedirect.com/topics/biochemistry-genetics-and-molecular-biology/fundulus-heteroclitus). Annotated and identified metabolite features were statistically analyzed using log normalization and pareto-scaling provided on metaboanalyst 5.0 (http://metaboanalyst.ca.) server. Pathway enrichment analysis was also performed using metaboanalyst 5.0.

***Analysis of Plasma cytokines using Bead Array Assay:***

The concentration of nineteen cytokines including pro and anti-inflammatory; TNF-α, IFN-α, IFN-β, IFN- γ, IL-1β, IL-17a, IL-18, IL-29 ( Pro-inflammatory) IL-2, IL-4, IL-6, IL-7, IL-8, IL-9, IL-10, IL-15, IL-21, IL-22, and IL-33 (Anti-inflammatory) were estimated using 25μl of plasma samples of all subjects (20 peripheral blood and 15 cord blood) using ProcartaPlex cytokine bead assay (ThermoFisher Scientific, Austria). A standard curve was plotted using the standards provided in the kit which corresponded to evaluate the respective cytokine values in samples. The lower limits of detection of the test for each cytokine is given in the supplementary table 1.

**Supplementary Table 1.** List of cytokines with their lower limit of detection.

| S.No. | Analytes | Lower Limits (pg/ml) |
| --- | --- | --- |
| 1 | TNF-α | 5.93 |
| 2 | IFN-α | 0.61 |
| 3 | IFN- β | 8.79 |
| 4 | IFN-γ | 17.92 |
| 5 | IL-1β | 2.56 |
| 6 | IL-17A | 3.1 |
| 7 | IL-18 | 18.63 |
| 8 | IL-2 | 8.18 |
| 9 | IL-4 | 15.09 |
| 10 | IL-6 | 10.55 |
| 11 | IL-7 | 0.9 |
| 12 | IL-8 | 2.76 |
| 12 | IL-9 | 9.74 |
| 13 | IL-10 | 2.29 |
| 14 | IL-15 | 3.54 |
| 15 | IL-17a | 3.1 |
| 15 | IL-18 | 18.63 |
| 16 | IL-21 | 9.3 |
| 17 | IL-22 | 21.24 |
| 18 | IL-29 | 59.4 |
| 19 | IL-33 | 22.85 |

***Whole blood immunophenotyping using high-dimensional flow cytometry analysis:***

Immunophenotyping of whole blood samples was carried out to characterize the monocytes, dendritic cells, T-cells, B cells, Tregs and exhaustion markers on them. In brief, 100-120µl of whole blood containing approximately, >1×10^5^ cells were lysed with FACS lyse solution (BD FACS^TM^ Lysing Solution cat no. #349202). Further, cells were washed twice with 1X PBS containing 0.5% BSA and 0.1% sodium azide. Cell pellets were incubated for 30 minutes at room temperature in the dark with specific antibodies against cell surface and intracellular markers. Antibodies were labelled with different fluorochromes and used for detection of monocytes, dendritic cells, T cells, and B cells ; HLA-DR, CD14, CD16, CD11C, CD123, CD11C, B cell; CD19, CD27, IgD, CD38, CD20, PDL1, CD152 (CTLA4), PD1, CD4, CD3, CD45RO, CD25, CCR7. All samples were processed in a single day in one batch to avoid batch variation. After incubation, the cells were washed and spin at 400g for 3 minutes and supernatant was discarded. 140 µl of cytofix and cytoperm (BD Biosciences, #554714) was added and then incubated for 10 min. after incubation, 500 µl of 1X perm wash (BD Biosciences, #554714) was added and centrifuged at 200g for 5 mins to remove supernatant. Similarly, samples were again washed with perm wash and finally cell pellet was incubated with intracellular antibodies for 20 min at room temperature. After incubation again cells were were washed with 1X PBS and a minimum of 100,000 events was acquired using BD FACSVerse™ flow Cytometer. For flow cytometric analysis, single-color compensation for each fluorochromes was performed. Data was analyzed using Flow Jo software version v10.8 (BD Biosciences).

***RNA isolation from placental tissue and Quantitative Real-time PCR***

Total RNA was extracted from the COVID-19 infected placenta (n=23) and healthy (n=7) using TRIzol reagent (Thermo Fisher) according to the manufacturer’s protocol. Isolated total RNA was quantified using NanoDrop 2000 (Thermo Scientifc) and 2ug of total RNA was used to synthesise complementary DNA (cDNA) using with High-capacity cDNA kit (ABI, Thermo Fisher Scientific, Waltham, MA, USA; #4368813). For analysing the quantitative expression of the genes, the cDNA prepared was mixed with SYBR Green Master Mix (Thermo Fisher, #A25742) and the reactions were carried out in a CFX-96 Biorad thermal cycler (Biorad, CA, USA).

**Primers List:**

| S.No. | Gene | Forward Primer (5’-3’) | Reverse Primer (5’-3’) |
| --- | --- | --- | --- |
| 1. | CD55 | TTTCCAGGACAACCAAGCATT | ACACGTGTGCCCAGATAGA |
| 2. | TNF- α | TGTAGCCCATGTTGTAGCAAAC | AGAGGACCTGGGAGTAGATGA |
| 3. | ESR-1 | CCAACCAGTGCACCATTGAT | GGTCTTTTCGTATCCCACCTTT |
| 4. | ESR-2 | GGCAGAGGACAGTAAAAGCA | GGACCACACAGCAGAAAGAT |
| 5. | GPER-1 | GTACTTCATCAACCTGGCGGTG | TCATCCAGGTGAGGAAGAAGACG |
| 6. | IFN- αR | GTCAGGCTGGTCTTGGACTN | GCCTCTTCAAAGACGCAAAN |

**Statistical analysis:**

Data was statistically analyzed using Prism (version 8.01; GraphPad). The comparison for continuous data is carried out by using the t-test/ one-way ANOVA/ Kruskal–Wallis test followed by probability adjustment by the Mann–Whitney test or by Bonferroni test post-hoc comparison as appropriate, and it is represented as mean ± SD. p-values <0.05 were considered as significant.

**Results:**

**Supplementary Table 2.** Demographic and Clinical characteristics of COVID-19 positive pregnant women with disease severity.

|  | ASYMPTOMATIC (n=82) | MILD (n=21) | MODERATE (n=9) | p value |
| --- | --- | --- | --- | --- |
| Age (years; median) | 25 (20-35) | 27.5 (21-35) | 25 (21-30) | 0.02 |
| Gestational age (months; median) | 38 (29-41) | 38 (34-39) | 39 (37-40) | 0.47 |
| Parity (median) | 1 (1-3) | 1 (1-3) | 1 | 0.42 |
| Normal vaginal delivery (%) | 67% | 58% | 66.70% | - |
| Caesarean section (%) | 33% | 42% | 33.30% | - |
| Apgar score at 5 min (median) | 50% (9,9,9) | 30% (9,9,9) | 100% (9,9,9) | - |
| Clinical parameters |  |  |  |  |
| Hb (mg/dL) | 10.8 (6.7-14.4) | 11 (6.8-13.3) | 12 (9-13.5) | 0.37 |
| Platelet count (Lakhs) | 1.64 (0.32-5.7) | 13.35 (0.28-2.8) | 2.8 (1.9-21.8) | 0.06 |
| TLC (µL) | 8945 (990-18690) | 10230 (1200-21200) | 12000 (8300-16300) | 0.1 |
| Total Bilirubin (mg/dL) | 0.6 (0.1-2.8) | 0.6 (0.2-1.7) | 0.4 (0.2-0.5) | 0.15 |
| AST (µ/L) | 31 (7-218) | 74 (12-337) | 38 (15-91 | 0.06 |
| ALT (µ/L) | 41 (11-275) | 565 (10-390) | 56 (23-116) | 0.29 |
| ALP (µ/L) | 221 (70-627) | 248.5 (29-689) | 236 (128-486) | 0.9 |
| SPO2 (%) | 98 (95-100) | 98 (88-100) | 99 (98-99) | 0.009 |
| D-Dimer (ng/mL) | 656 (125-5000) | 502 (199-2235) | 810 (367-910) | 0.43 |
| PT (sec) | 11.15 (8.6-72) | 11.1 (10.2-17.5) | 10.2 (9.6-11.2) | 0.63 |
| INR (sec) | 0.9 (0.7-4.8) | 0.9 (0.8-1.4) | 0.9 (0.7- 0.93) | 0.64 |
| Urea (mg/dL) | 16.5 (7-37) | 17 (0.6-35) | 34 (29-32) | 1.5 |
| Creatinine (mg/dL) | 0.475 (0.3-10.4) | 0.575 (0.4-11.8) | 0.6 (0.5-0.7) | 0.15 |
| Na (mmol/L) | 136 (128-144) | 138 (134-147) | 140 (132- 141) | 0.001 |
| K (mmol/L) | 4.1 (3.3-7.5) | 4.3 (3.3-5.3) | 4.8 (4.7- 5.1) | 0.012 |

***Supplementary Figure 1***

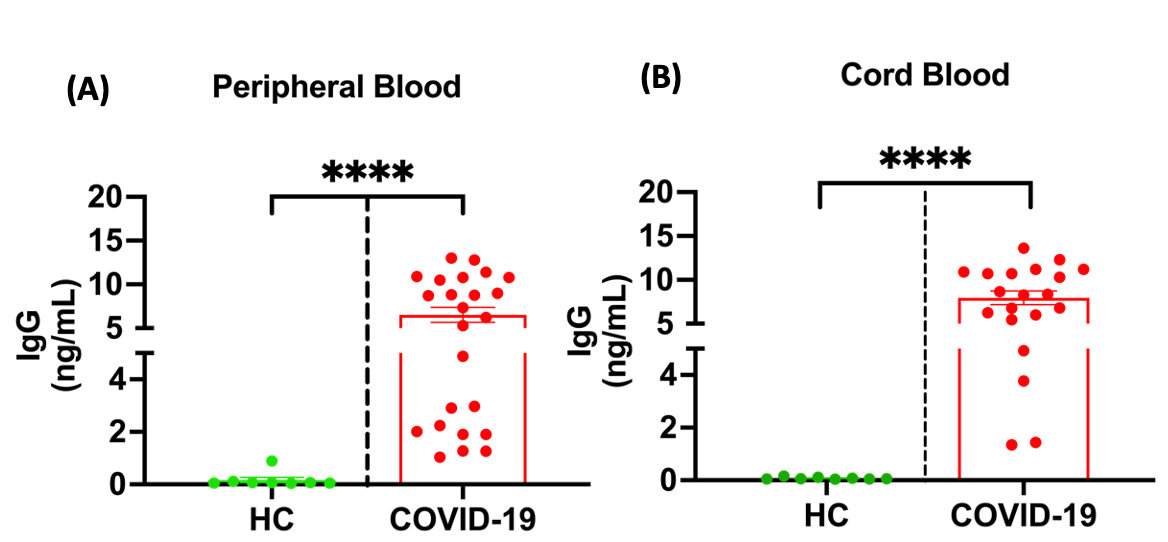

**Supplementary Figure 1. IgG titers in COVID-19 positive pregnant women peripheral and cord blood (A)** Plasma concentrations of IgG in the maternal blood healthy, [n = 8] and COVID-19 positive [n=24] **(B)** Plasma concentrations of IgG in cord blood [n = 8 healthy, 20 COVID-19 positive. Significance differences is based on p<0.05.

***Supplementary Figure 2***

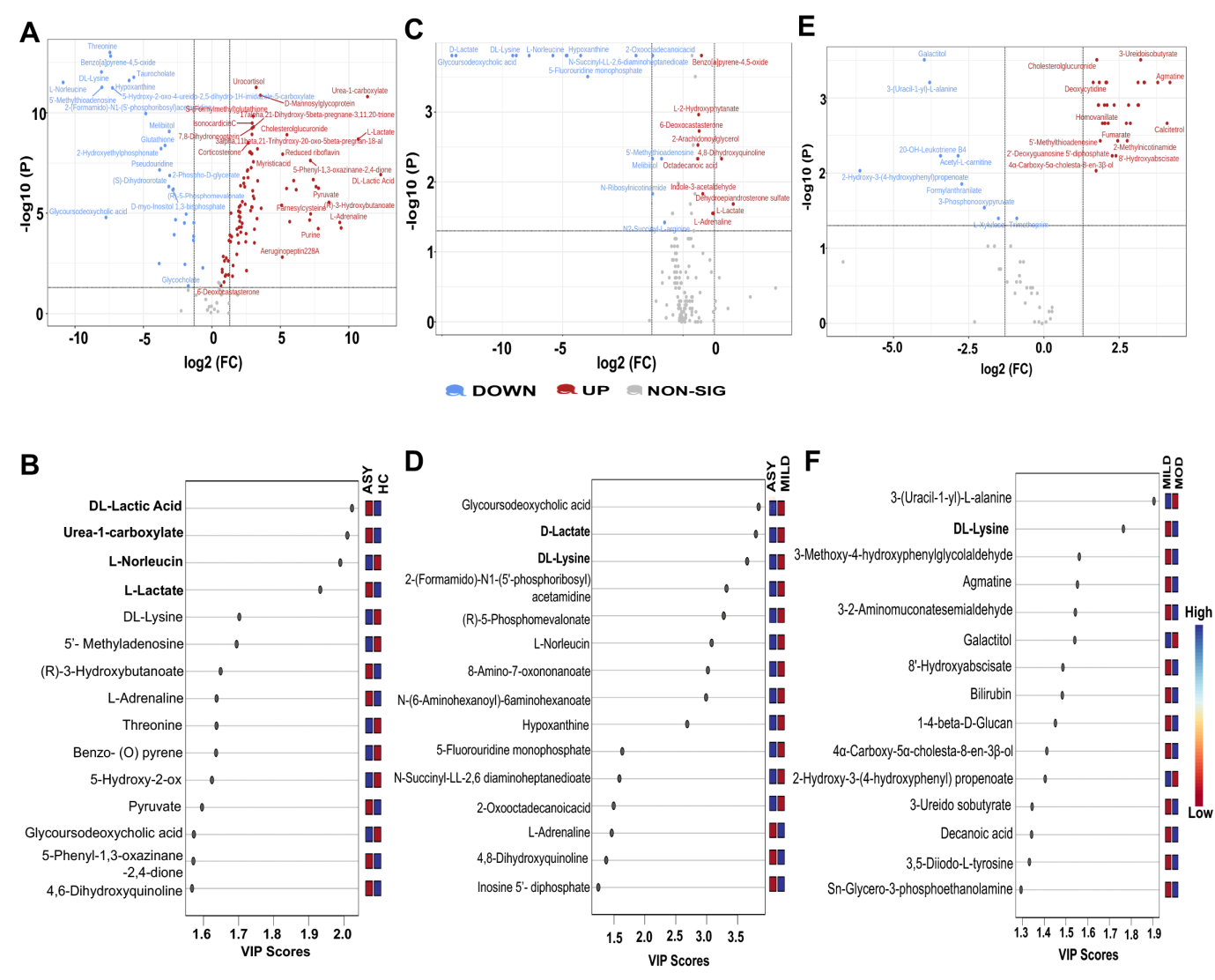

**Supplementary Figure 2. Differentially expressed metabolites with increased disease severity. (A-B; HC vs. ASY) A.** The volcano plot shows the differential expression of metabolites; in asymptomatic COVID-19 positive compared to healthy. Red and blue dots represent metabolites that are significantly up regulated (fold-change > 2 and p-value < 0.05), and down regulated. **B.** Variable importance in projection (VIP) plot represented the separation of metabolite profiles between COVID-19 infected pregnant women (red) and healthy (blue). **(C-D; ASY vs. MILD) C.** The volcano plot shows the differential expression of metabolites; in mild COVID-19 positive compared to asymptomatic group. Red and blue dots represent metabolites that are significantly up regulated (fold-change > 2 and p-value < 0.05), and down regulated. **D.** VIP plot represented the separation of metabolite profiles between infected pregnant women (red) and healthy (blue). **(E-F; MILD vs. MOD) E.** The volcano plot shows the differential expression of metabolites; in moderate COVID-19 positive compared to mild group. Red and blue dots represent metabolites that are significantly up regulated (fold-change > 2 and p-value < 0.05), and down regulated. **F.** VIP plot represented the separation of metabolite profiles between Mild infected pregnant women (red) and moderate (blue).

**Supplementary Table 3. Alteration in metabolic pathways in COVID-19 positive pregnant women with disease severity**

|  | **Metabolic pathways** | |
| --- | --- | --- |
|  | **Upregulated** | **Downregulated** |
| **Healthy vs. Asymptomatic** | - Valine, leucine, isoleucine - Taurine, hypotaurine metabolism - Hypotaurine metabolism - **Glycine, serine and threonine metabolism** | - One carbon pool by folate - **Riboflavin** - **Alanine, aspartate and glutamate** - **Tyrosine** |
| **Asymptomatic vs. Mild** | - Nicotinate and nicotinamide metabolism - Pyruvate metabolism | - **Riboflavin** - Biotin metabolism |
| **Mild vs. Moderate** | - Pentose and glucuronate interconversions - **Glycine, serine and threonine metabolism** - Galactose metabolism | - **Riboflavin** - **Tyrosine** - **Alanine, aspartate and glutamate** |

***Supplementary Figure 3***

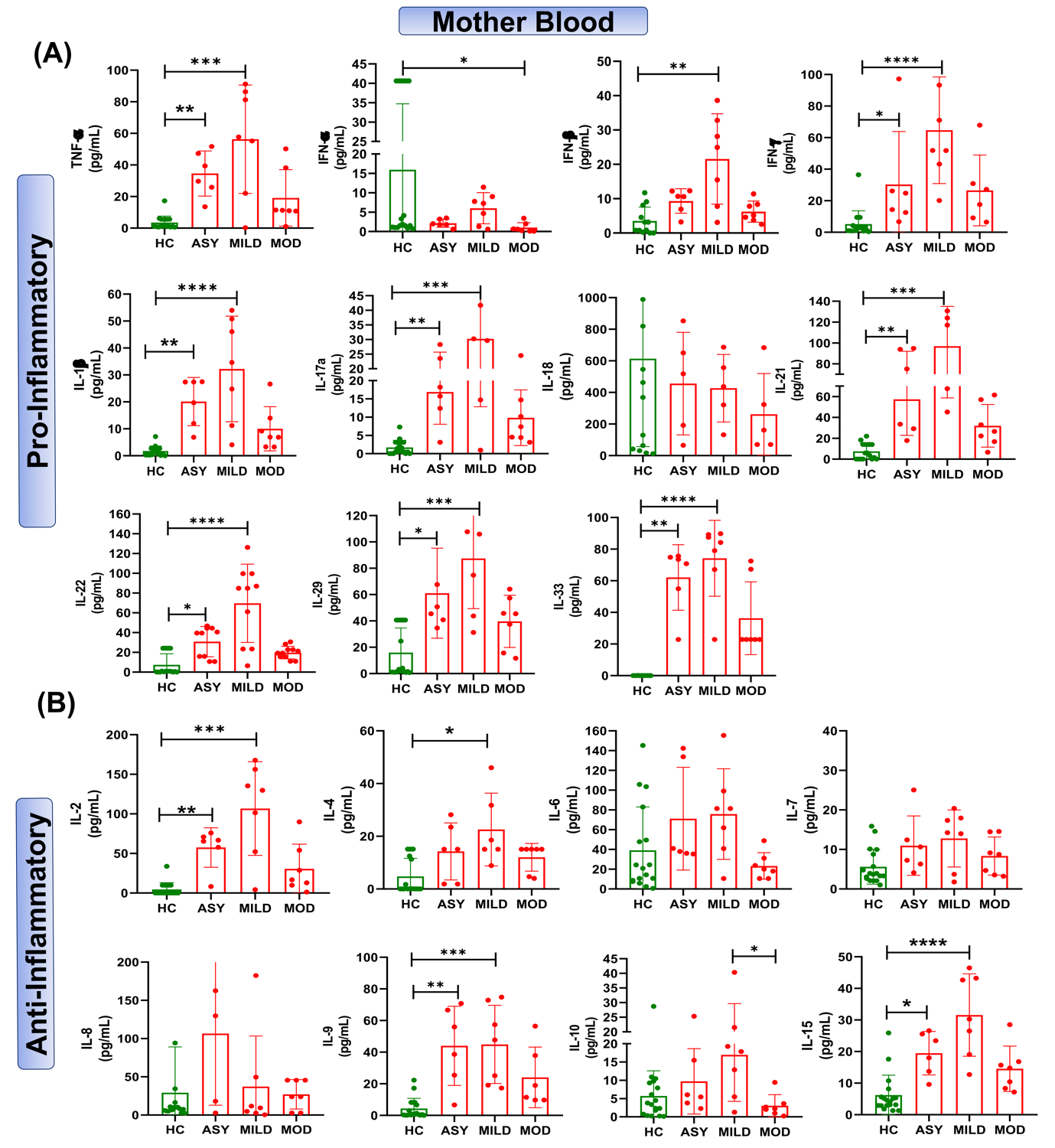

**Supplementary Figure 3.** Cytokine levels in COVID-19 positive pregnant women based on disease severity (asymptomatic (n=6), mild (n=7) and moderate (n=7)). **(A)** Pro-inflammatory cytokines **(B)** Anti-inflammatory cytokines. Data shown as boxplots and presented as mean ± standard deviation (SD). Data was analyzed using unpaired t-test with Mann-Whitney correction and one way ANOVA. Significance levels are indicated as *p<0.05, **p<0.01, ***p<0.001, and ****p<0.001.

***Supplementary Figure 4***

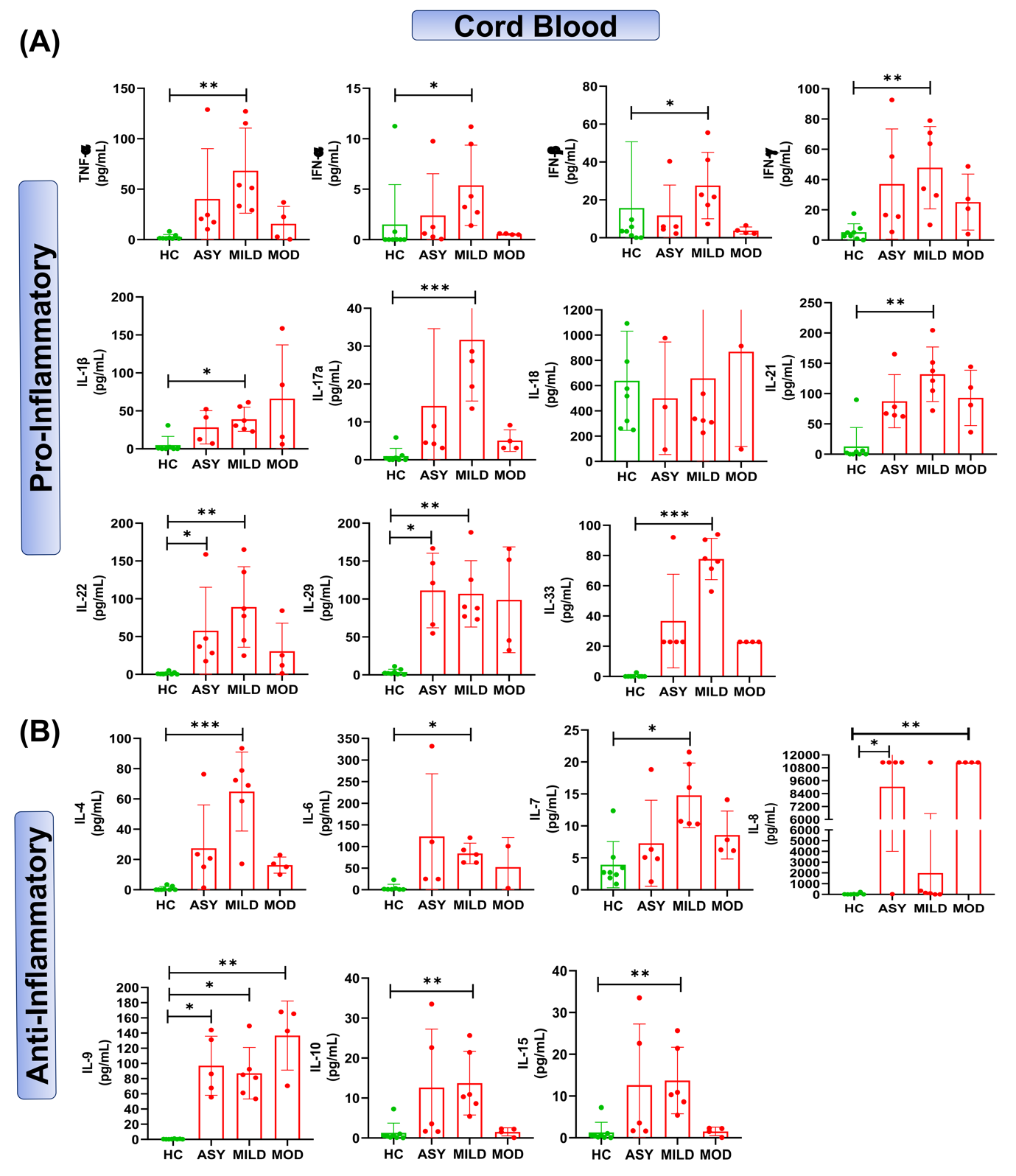

**Supplementary Figure 4.** Cytokine concentrations in the cord blood of COVID-19 positive asymptomatic (n=5), mild (n=6), moderate (n=4) and healthy control (n=18) women. Data are shown as boxplots and presented as mean ± standard deviation (SD). Data was analyzed using unpaired t-test with Mann-Whitney correction and one way ANOVA. Significance levels are indicated as *p<0.05, **p<0.01, and ***p<0.001.

**Supplementary Figure 5**

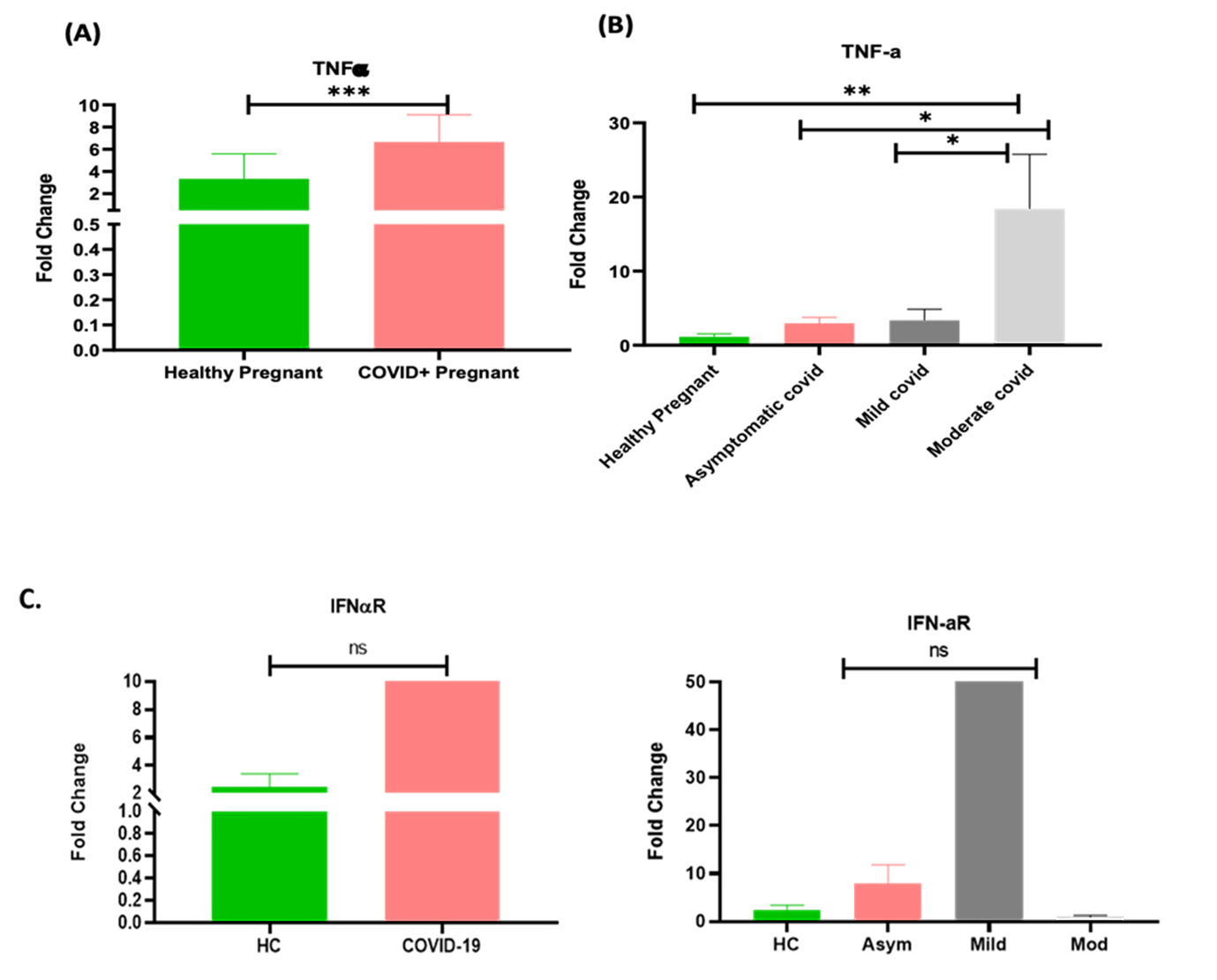
**Supplementary Figure 5. Expression of TNF-α and IFN-αR in the placenta of COVID-19 infected mothers (A)** TNF-**α** expression in healthy controls (n=7) and COVID-19 placenta (n=23). **(B)** TNF-**α** expression asymptomatic (n=13), mild (n=7) and moderate (n=3). **(C)** IFN-αR expression in expression in healthy and COVID-19 infected placentas.

***Supplementary Figure 6***

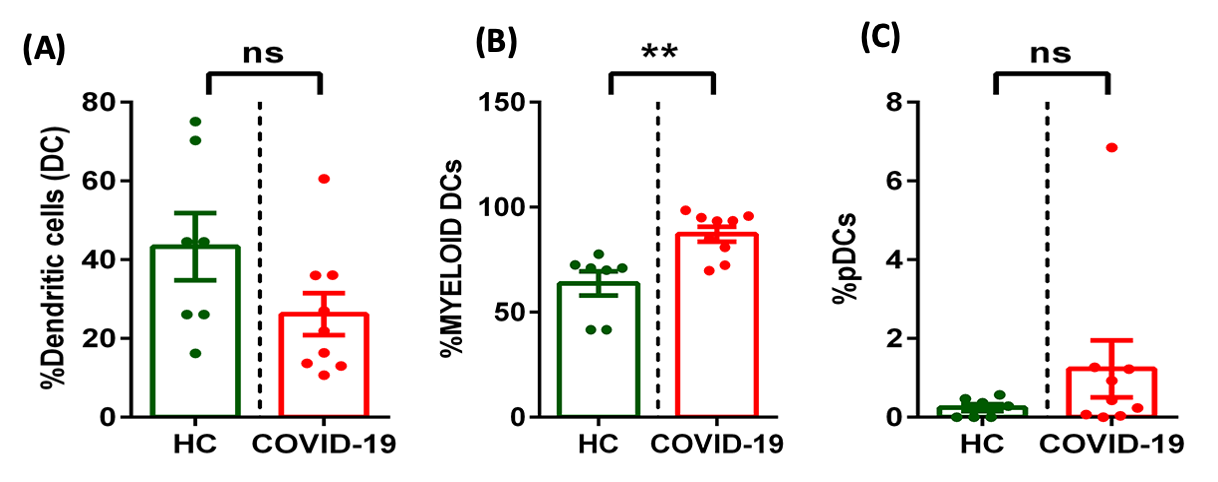

**Supplementary Figure 6. Immunophenotyping of dendritic cells in the pregnant mothers (A)** Frequency of absolute dendritic cells in healthy pregnant control (n=7) and SARS-CoV-2 (+ve) (n=9) **(B)** Frequency of myeloid DCs (mDCs) **(C)** Frequency of plasmacytoid dendritic cells(pDCs). The data are presented as mean ± standard deviation (SD) and analyzed using unpaired t-test with Mann-Whitney correction. Significance levels are indicated as **p<0.01.

***Supplementary Figure 7***

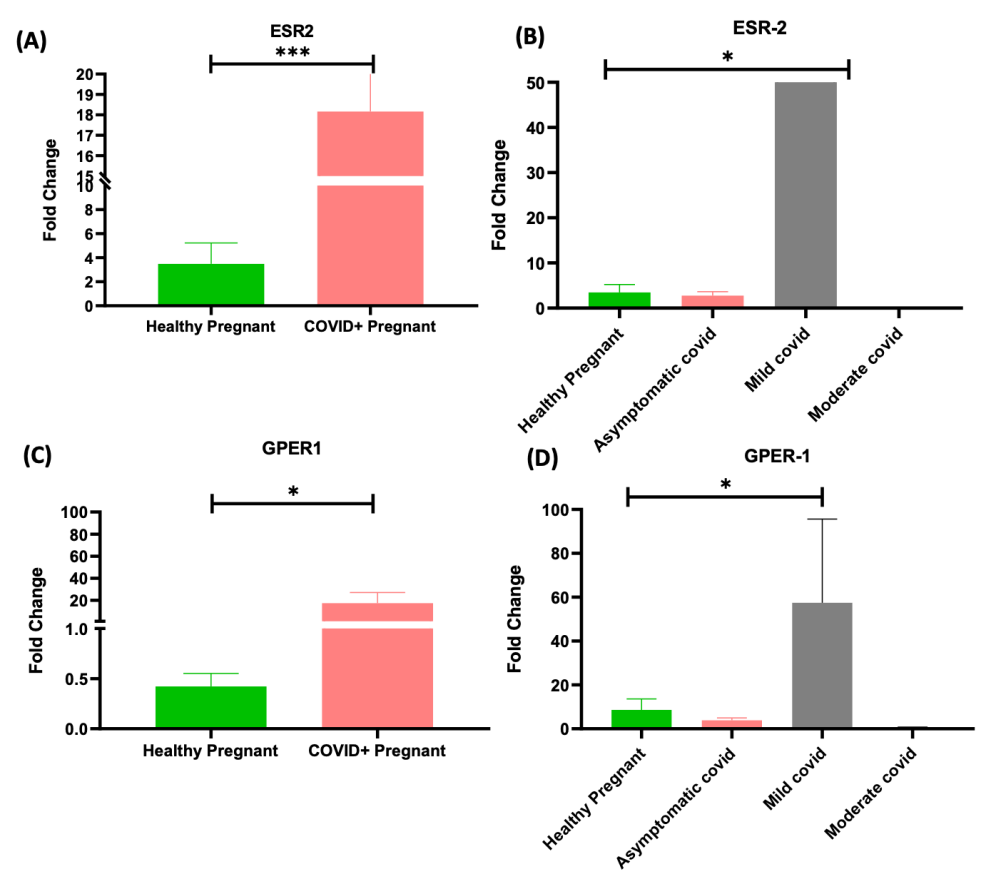

**Supplementary Figure 7. Expression of estrogen receptors in placenta of COVID-19 infected mothers (A)** Estrogen receptor beta **(**ESR2) expression in healthy controls (n=7) and COVID-19 (n=23). **(B)** Estrogen receptor beta **(**ESR2) expression in healthy (n=7), asymptomatic (n=13), mild (n=7) and moderate (n=3). **(C)** G-protein coupled estrogen receptor beta **(**GPER-1) expression in healthy controls (n=7) and COVID-19 (n=23). **(D)** G- protein coupled estrogen receptor **(**GPER-1) expression in healthy controls (n=7), asymptomatic (n=13), mild (n=7) and moderate (n=3). The data is presented as mean ± standard deviation (SD) and analyzed using unpaired t-test with Mann-Whitney correction and one way ANOVA. Significance levels are indicated as *p<0.05 and ***p<0.001.
